# Decoding sequence determinants of gene expression in diverse cellular and disease states

**DOI:** 10.1101/2024.10.09.617507

**Authors:** Avantika Lal, Alexander Karollus, Laura Gunsalus, David Garfield, Surag Nair, Alex M Tseng, M Grace Gordon, John Blischak, Bryce van de Geijn, Tushar Bhangale, Jenna L Collier, Nathaniel Diamant, Tommaso Biancalani, Hector Corrada Bravo, Gabriele Scalia, Gokcen Eraslan

## Abstract

Sequence-to-function models that predict gene expression from genomic DNA sequence have proven valuable for many biological tasks, including understanding *cis*-regulatory syntax and interpreting non-coding genetic variants. However, current state-of-the-art models have been trained largely on bulk expression profiles from healthy tissues or cell lines, and have not learned the properties of precise cell types and states that are captured in large-scale single-cell transcriptomic datasets. Thus, they lack the ability to perform these tasks at the resolution of specific cell types or states across diverse tissue and disease contexts. To address this gap, we present Decima, a model that predicts the cell type- and condition- specific expression of a gene from its surrounding DNA sequence. Decima is trained on single-cell or single-nucleus RNA sequencing data from over 22 million cells, and successfully predicts the cell type-specific expression of unseen genes based on their sequence alone. Here, we demonstrate Decima’s ability to reveal the *cis*-regulatory mechanisms driving cell type-specific gene expression and its changes in disease, to predict non-coding variant effects at cell type resolution, and to design regulatory DNA elements with precisely tuned, context-specific functions.

## 1. Introduction

Sequence-to-function deep learning models that predict gene expression from genomic DNA sequences have proven valuable for many biological tasks, including understanding *cis*-regulatory mechanisms and interpreting non-coding genetic variants (Zhou et al. 2018; Trevino et al. 2021; K. M. Chen et al. 2022; Eraslan et al. 2019; Sasse, Chikina, and Mostafavi 2024; Agarwal and Shendure 2020; Avsec, Weilert, et al. 2021). Recently, modeling the genomic context up to hundreds of kilobases has enabled effective prediction of bulk cap analysis of gene expression (CAGE-seq) (Avsec, Agarwal, et al. 2021) and RNA-seq (Linder et al. 2023) profiles. However, current state-of-the-art models have been trained largely on bulk expression profiles from healthy tissues or cell lines (K. M. Chen et al. 2022; Avsec, Agarwal, et al. 2021; Linder et al. 2023). Thus, they cannot predict expression nor reveal regulatory mechanisms that act in specific cell types. Moreover, due to training primarily on samples from healthy donors, they cannot directly model pathological expression changes occurring in the context of disease. Therefore they fall short of learning the extensive biological information embedded in the rapidly accumulating single-cell or single-nucleus RNA sequencing (sc/snRNA-seq) datasets.

In addition, studying regulatory mechanisms using scRNA-seq data has been challenging but holds great potential. Current approaches typically rely on integrating single-cell ATAC-seq (scATAC) data, or analyzing the expression of transcription factors (TFs) or their known target genes, or analyzing the co-expression of TFs and targets. While effective, these approaches are limited by the availability of matched scATAC-seq or prior knowledge of TF-target relationships (Holland et al. 2020; Badia-i-Mompel et al. 2023). However, the genomic sequence itself offers a rich and largely untapped data modality for uncovering regulatory mechanisms. By focusing on the sequence, we can move beyond the constraints of known TF-target pairs and explore the regulatory landscape of biological settings for which chromatin accessibility data are unavailable, opening the door to a more comprehensive understanding of gene regulation across diverse cell types and conditions.

As a result, there is a need for sequence models that can uncover complex regulatory mechanisms at play within cell populations across different conditions. This gap has been widely recognized, leading to recent attempts to predict gene expression from sequence in single cells (Schwessinger et al. 2023; Jiaqi Li et al. 2022; Hingerl et al. 2024). However, such an approach is difficult to scale to atlas-level single-cell datasets that frequently include millions of cells, and is consequently limited to analyses of single datasets comprising a small number of cell types or states.

Here we present **Decima**, a model that predicts the cell type- and condition-specific expression of a gene from its surrounding genomic sequence. Decima is trained on sc/snRNA-seq count data aggregated at pseudobulk resolution, allowing it to learn from a massive training corpus comprising data from over 22 million cells, and representing 201 distinct cell types in 271 tissues and 82 diseases. We demonstrate Decima’s ability to accurately predict pseudobulk gene expression, including highly cell type-specific genes, across studies and conditions. Additionally, Decima identifies potential regulatory elements driving cell type- and cell state-specific gene expression, as well as tissue residency and disease-associated expression changes. We also show that Decima can predict the effects of non-coding genetic variants in individual cell types and present a proof-of-concept example where Decima facilitates the design of disease-biased regulatory elements.

## 2. Results

### 2.1 Decima predicts pseudobulk gene expression from DNA sequence

We obtained annotated sc/snRNA-seq atlases from SCimilarity (Heimberg et al. 2023), the human brain atlas (X. Chen et al. 2024), skin atlas (Fiskin et al. 2023), and adult human retina atlas (Jin Li et al. 2023) (**Supplementary Table 1**). After processing and filtering **(Methods)**, our training corpus comprised data from over 22 million cells. We summed the RNA-seq counts from all cells belonging to the same combination of cell type, tissue, disease state, and study, resulting in a final count matrix consisting of 8,856 pseudobulk expression vectors and 18,457 genes. This matrix includes pseudobulks representing 201 distinct cell types in 271 tissues and 82 diseases; there are 6,481 unique combinations of cell type, tissue, and disease state, and 4,478 unique combinations of cell type and tissue annotations. We normalized the counts in each pseudobulk to CPM (counts per million) values and applied a log(1+x) transformation for variance stabilization.

Each example seen by Decima is focused on a single gene. For each gene, we created an input window containing 524,288 bp of genomic sequence surrounding the gene. This input window contained the gene transcription start site (TSS) and a minimum of 163,840 bp upstream of the TSS, with the remaining sequence covering the gene body and downstream regions (**Supplementary Fig. 1**). In addition to the one-hot encoded DNA sequence in the input window, we also supplied the model with a binary mask representing the location of the gene body in the input window **(Fig. 1A)**.

**Fig. 1.**
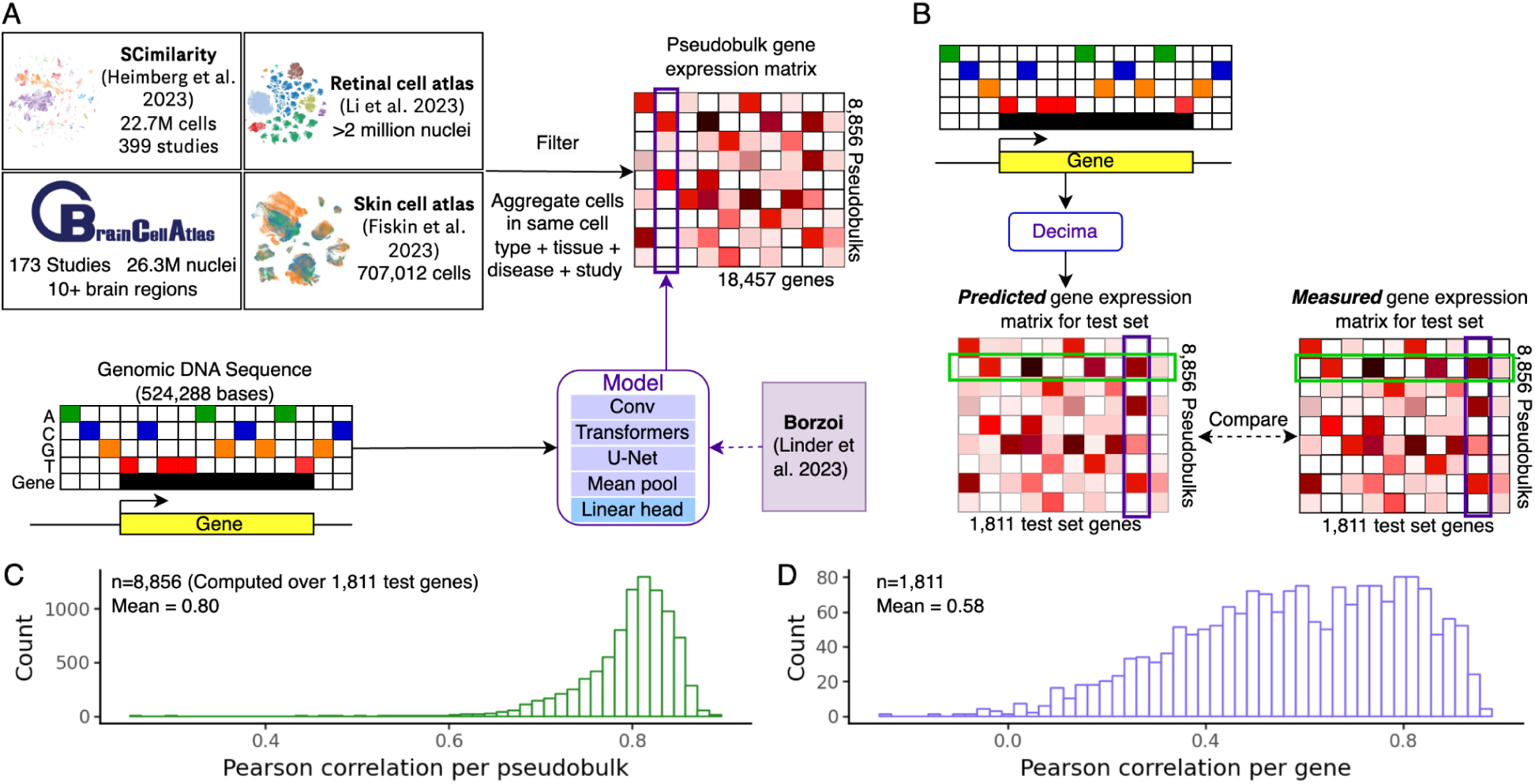
**A)** Schematic of the Decima model. Single-cell datasets were combined, filtered, and aggregated to create a pseudobulk gene expression matrix. Each row of this matrix corresponds to a unique combination of cell type, tissue, disease, and study, and each column corresponds to a single gene. The model takes as input the DNA sequence surrounding a single gene and predicts the corresponding column of the expression matrix. **B)** Schematic showing the evaluation of the trained Decima model on 1,811 test set genes. **C)** Histogram of Pearson correlation coefficients between the measured and predicted expression vectors for each of the 8,856 pseudobulks (rows of the matrix in B), over 1,811 test set genes. **D)** Histogram of Pearson correlation coefficients between the measured and predicted per-gene expression vectors (columns of the matrix in B) for each of the 1,811 test set genes.

We initialized Decima with the recent Borzoi model (Linder et al. 2023), which has been trained to predict bulk expression profiles (RNA-seq and CAGE-seq) as well as epigenomic profiles (DNase-seq, ATAC-seq, and ChIP-seq) from the genome sequence. This allows the model to benefit from learned epigenetic regulatory mechanisms, which are more easily learned from these assays than from scRNA-seq data, as well as from learned rules related to gene expression at bulk resolution. We replaced the final layer of Borzoi with a global mean pooling layer along the length axis, followed by a linear layer that outputs a vector of 8,856 values for each gene, corresponding to the predicted log(CPM+1) values in each pseudobulk **(Fig. 1A)**. We split the 18,457 genes in the pseudobulk matrix into groups for training, validation, and testing based on their overlap with the genomic regions used to train Borzoi, such that the sequences in our test set were also in the test set of Borzoi.

We trained the entire model using a novel two-component loss function comprising a Poisson loss on the total expression of the gene across all pseudobulks, and a multinomial loss along the pseudobulk axis. This is based on the two-component loss introduced in (Avsec, Weilert, et al. 2021) and adapted for Borzoi (Linder et al. 2023); however, in these studies, the multinomial loss was calculated across sequence positions, encouraging the model to focus on between-region differences in biological activity. In contrast, our loss function encourages the model to correctly predict differences in expression between pseudobulks for a single gene.

We evaluated Decima by predicting the expression of 1,811 held-out genes in all pseudobulks **(Fig. 1B; Supplementary File 1)**. As Borzoi consists of an ensemble of 4 replicate models (https://github.com/calico/borzoi/blob/main/README.md), we trained 4 Decima models, each initialized with a different replicate, and averaged their predictions to increase robustness. Considering only held-out genes, the mean Pearson correlation between the measured and predicted expression vectors for each pseudobulk was 0.80, indicating that the model performs well at predicting differences in expression between unseen genes in the same condition **(Fig. 1C)**. This performance was largely consistent across the wide variety of datasets, cell types, and disease states included in the dataset **(Supplementary Fig. 2-5)**. Further, the mean correlation between the measured and predicted expression vectors for each held-out gene was 0.58, indicating that the model can also predict differences in expression of the same gene between different populations of cells **(Fig. 1D)**. This analysis thus validated that Decima successfully learned determinants of both between-gene and between-cell type expression changes.

### 2.2 Decima learns sequence determinants of cell type specificity

We tested whether Decima could successfully predict cell type-specific patterns of gene expression from sequence alone. We first identified test set genes specific to each cell type in the dataset based on their z-score normalized expression values **(Methods; Fig. 2A)**. Specifically, we labeled a gene as “specific” to a cell type if its average z-score in that cell type based on its measured expression was >=1. For each cell type, we tested whether the z-scores based on Decima’s predicted expression matrix could be used to classify the specific and nonspecific genes for that cell type (Eraslan et al. 2022) (**Fig. 2A**). We obtained an average AUROC of 0.82 across cell types **(Fig. 2B)**. This analysis suggests that Decima has learned sequence features that determine the expression patterns of even cell type-specific genes.

**Fig. 2.**
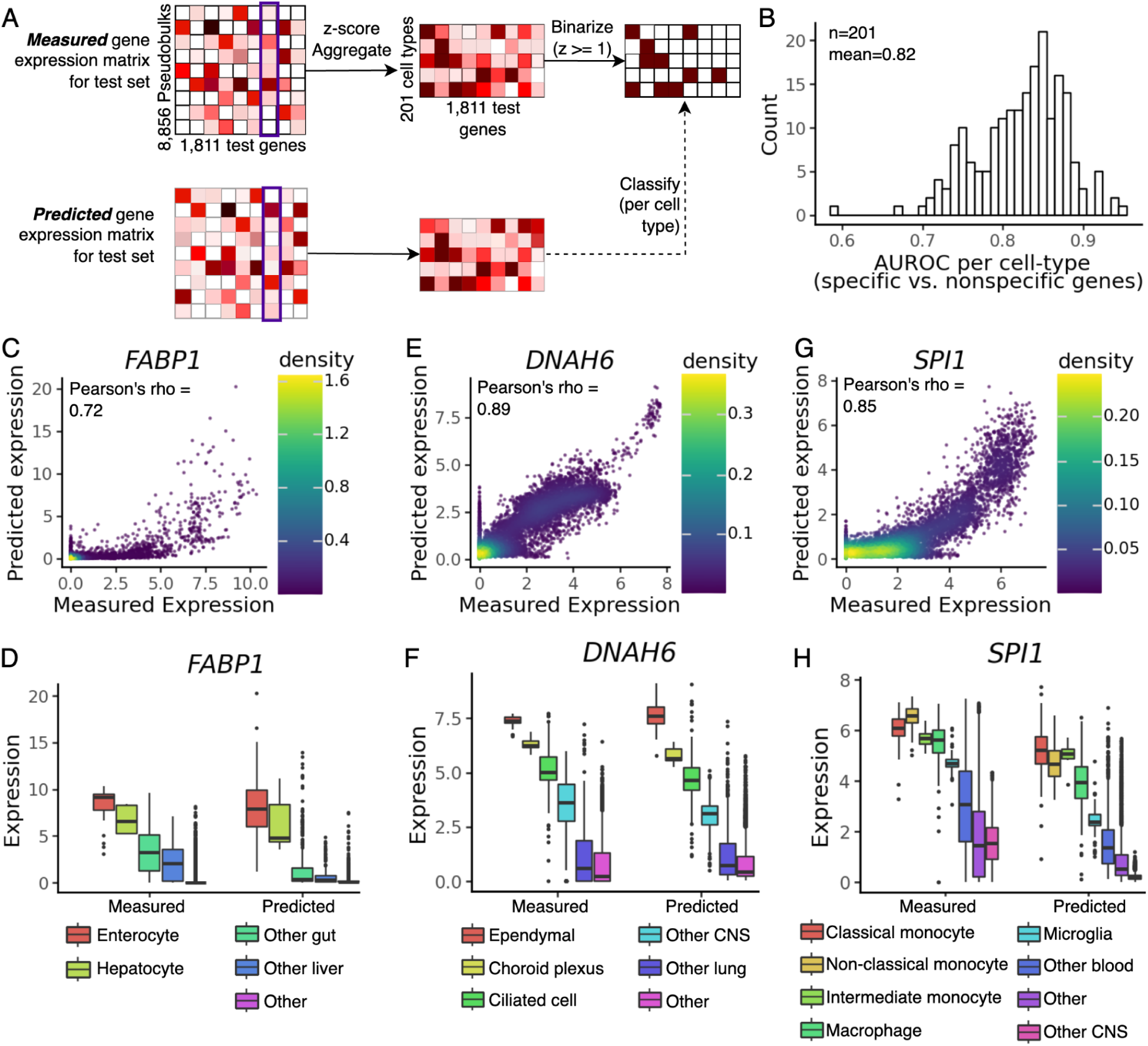
**A)** Schematic illustrating the approach used to evaluate Decima’s performance on identification of cell type-specific genes. **B)** Histogram showing the Area Under the Receiver Operator Characteristic (AUROC) for classification of specific vs. nonspecific genes in each cell type based on Decima’s predictions in the same cell type. **C)** Scatter plot showing the measured and predicted expression (log (CPM+1)) of the FABP1 gene in all pseudobulks. **D)** Boxplots showing the measured and predicted expression of the FABP1 gene in pseudobulks representing enterocytes and hepatocytes, compared to pseudobulks representing other cell types in the gut, other cell types in the liver, and all remaining cell types. **E)** Scatter plot showing the measured and predicted expression of the DNAH6 gene in all pseudobulks. **F)** Boxplots showing the expression of the DNAH6 gene in pseudobulks representing ependymal cells, choroid plexus cells, ciliated cells, other cell types in the central nervous system, other cell types in the lung, and all remaining cell types. **G)** Scatter plot showing the true and predicted expression of the SPI1 gene in all pseudobulks. **H)** Boxplots showing the expression of the SPI1 gene in pseudobulks representing monocytes, macrophages, microglia, other cell types in the blood, other cell types in the CNS, and all remaining cell types. In all boxplots, the lower and upper hinges correspond to the first and third quartiles, whiskers extend to 1.5 * IQR (inter-quartile range), and the remaining points are plotted individually.

We verified this by examining Decima’s predictions for individual genes with highly specific expression in diverse cell types. For example, the *FABP1* gene encodes a protein critical for the uptake and intracellular transport of fatty acids, and is highly expressed in the enterocytes of the intestinal lining and hepatocytes in the liver. Decima correctly predicts its elevated expression in both these cell types, distinguishing them from other cell types in the same tissues (**Fig. 2C, D**). Similarly, Decima correctly predicts the expression of *DNAH6*, a gene that encodes a component of cilia and is consequently highly expressed in the ependymal and choroid plexus cells of the brain and in the ciliated cells of the lung (**Fig. 2E, F**); and *SPI1*, which encodes a transcriptional regulator of myeloid cell development and is highly expressed in cells of the myeloid lineage (**Fig. 2G, H**).

We next asked how Decima distinguishes genomic features relevant to expression, including intron/exon boundaries and *cis*-regulatory elements. For each gene in the test set, we calculated Decima’s attributions across all nucleotides in the input genomic interval, using the Input x Gradient method (Shrikumar, Greenside, and Kundaje 2017). Gradients were calculated with respect to the average expression of the gene across all pseudobulks where the gene was strongly expressed (log(CPM + 1) > 1.5). We used the absolute value of these attributions, averaged across a genomic region, as a measure of the importance of that region to the predicted expression of the gene. On average, the magnitude of the attributions was highest within the gene promoter, followed by the exon/intron junctions and exons of the gene. In comparison, introns of the same gene had lower attributions. However, intronic cis-regulatory elements (CREs) had higher attributions than the rest of the intron **(Fig. 3A)**. Outside the gene body, attribution magnitude decayed with distance from the gene **(Fig. 3B)**. However, Decima assigned higher attributions to nucleotides within annotated cis-regulatory elements (ENCODE Project Consortium et al. 2020), even at elements far away (> 100 kb) from the predicted gene. This indicates that the model has learned to distinguish various classes of regulatory sequence elements, and uses these to inform its predictions.

**Fig. 3.**
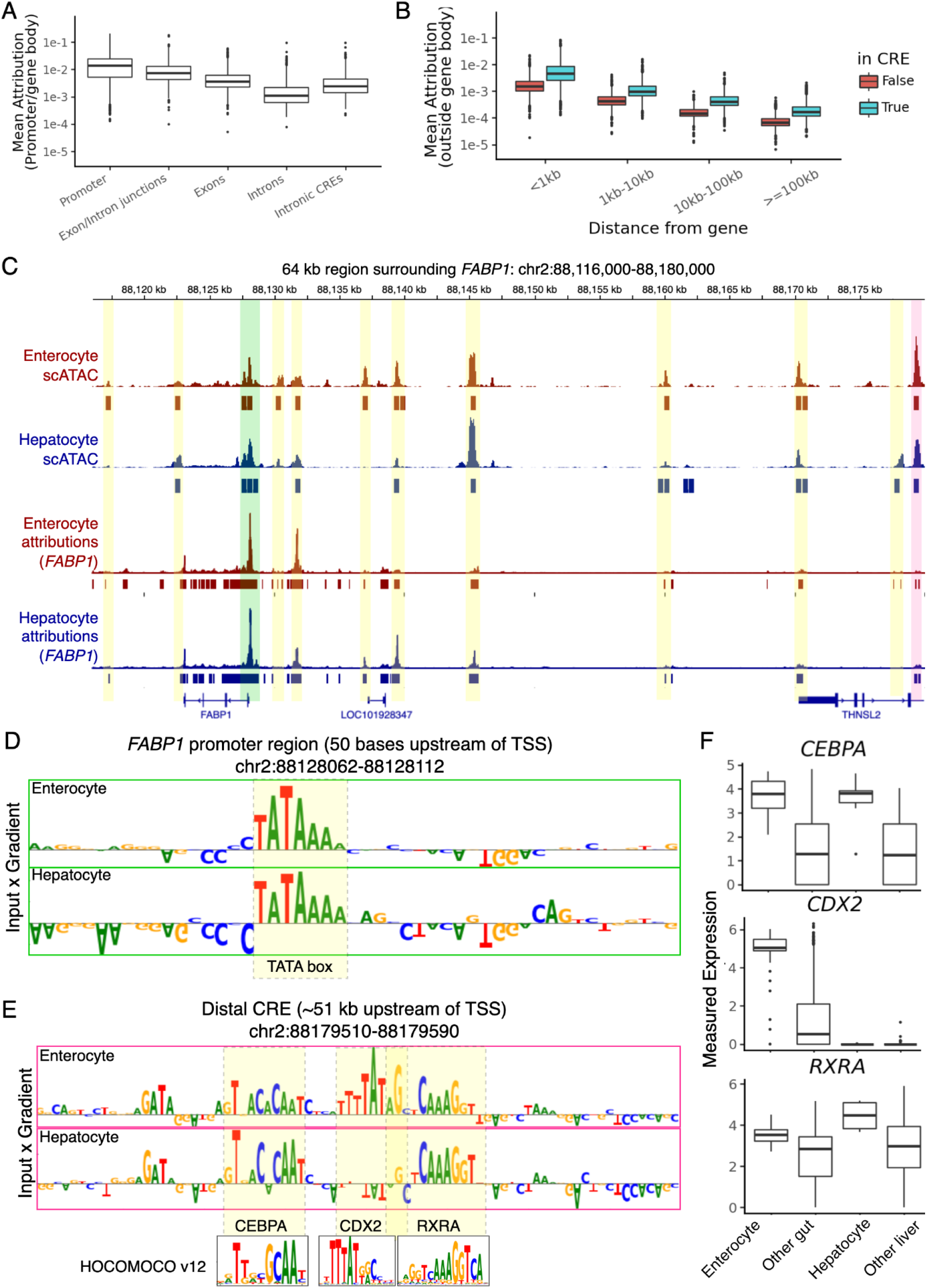
**A)** Per-nucleotide attribution scores averaged across different sequence elements in the gene body and promoter, for all test set genes. Values plotted represent the absolute value of the input x gradient score, averaged across all nucleotides of the corresponding type of element for each gene. **B)** Average per-nucleotide attribution scores within different distance ranges from the gene body, for all test set genes. Within each distance range, nucleotides within annotated cis-regulatory elements (CREs) are separated from other nucleotides. Values plotted represent the absolute value of the input x gradient score, averaged across all nucleotides in the corresponding distance range from each gene **C)** Upper: Pseudobulk scATAC-seq coverage (K. Zhang et al. 2021) over a 64 kb genomic region surrounding the FABP1 gene. Coverage tracks and peaks are shown for enterocytes (red) and hepatocytes (blue). Lower: Smoothed per-nucleotide attribution scores for predicted FABP1 expression over the same genomic region in enterocytes (red) and hepatocytes (blue). Values plotted represent the absolute value of the input x gradient score, smoothed with a gaussian filter of width 11 bp. scATAC-seq peaks are highlighted. **D)** Per-nucleotide input x gradient attribution scores over the 50 nucleotides upstream of the FABP1 transcription start site, in enterocytes (upper) and hepatocytes (lower). A TATA box is highlighted in yellow. **E)** Per-nucleotide input x gradient attribution scores over a distal CRE (also highlighted in pink in panel C), in enterocytes (upper) and hepatocytes (lower). Significant matches to HOCOMOCO v12 motifs are highlighted in yellow. **F)** Box plots showing the measured expression of the CEBPA, CDX2 and RXRA genes in pseudobulks representing enterocytes and hepatocytes, compared to pseudobulks representing other cell types in the gut and liver.

To investigate whether Decima could identify sequence features driving cell type-specific gene expression, we focused on its attributions in the genomic interval surrounding the previously mentioned *FABP1* gene **(Fig. 2D)**. In both enterocytes and hepatocytes, Decima’s attributions for this gene highlighted several regions corresponding to accessible chromatin (K. Zhang et al. 2021), including regions as far as > 50 kb from the transcription start site **(Fig. 3C)**.

Closer examination of regions with the highest attribution revealed known regulatory motifs, such as a TATA box in the *FABP1* promoter, which had a positive contribution to *FABP1* expression in both cell types **(Fig. 3D)**. In a distal CRE over 50 kb from the TSS which is accessible in both cell types **(Fig. 3C)**, Decima’s attributions highlighted C/EBP, CDX, and RXR transcription factor (TF) binding motifs in enterocytes; however, only the C/EBP and RXR motifs showed high attributions in hepatocytes **(Fig. 3E)**. This may be explained by the elevated expression of the *CEBPA* and *RXRA* genes encoding these TFs in both enterocytes and hepatocytes, whereas the *CDX2* gene shows elevated expression only in enterocytes **(Fig. 3F)**, consistent with literature (Grainger, Hryniuk, and Lohnes 2013). This example demonstrates that Decima has learned not only general sequence features such as gene and exon/intron boundaries, but also motifs with cell type-specific regulatory activity, and that Decima’s attributions can identify the sequence features driving cell type-specific expression patterns.

### 2.3 Decima’s attributions reveal drivers of cell type identity, cell state and tissue residency

Given that Decima can predict the cell type specificity of individual genes, we asked whether examining the sequence features with high attribution scores for all genes specific to a cell type would reveal consistent motifs for transcription factors underlying cell type identity. The procedure for this analysis is illustrated in **Fig. 4A**. We calculated the predicted log fold change in gene expression between the cell type of interest (‘positive’) and other ‘negative’ cell types in the same tissue. We identified genes with the highest cell type-specificity in the ‘positive’ cell type, and for each of these genes, used the Input x gradient method to compute the attribution of its predicted log fold change with respect to each nucleotide in the input DNA sequence. We refer to this score as the “differential attribution” for each nucleotide. We then applied TF-MoDISco (Shrikumar et al. 2018) to cluster these differential attributions and identify motifs with consistently high contributions. We quantified the contribution of each motif with an aggregate contribution score representing its contribution to cell type- specific gene expression in the ‘positive’ cell type **(Methods)**.

**Fig. 4.**
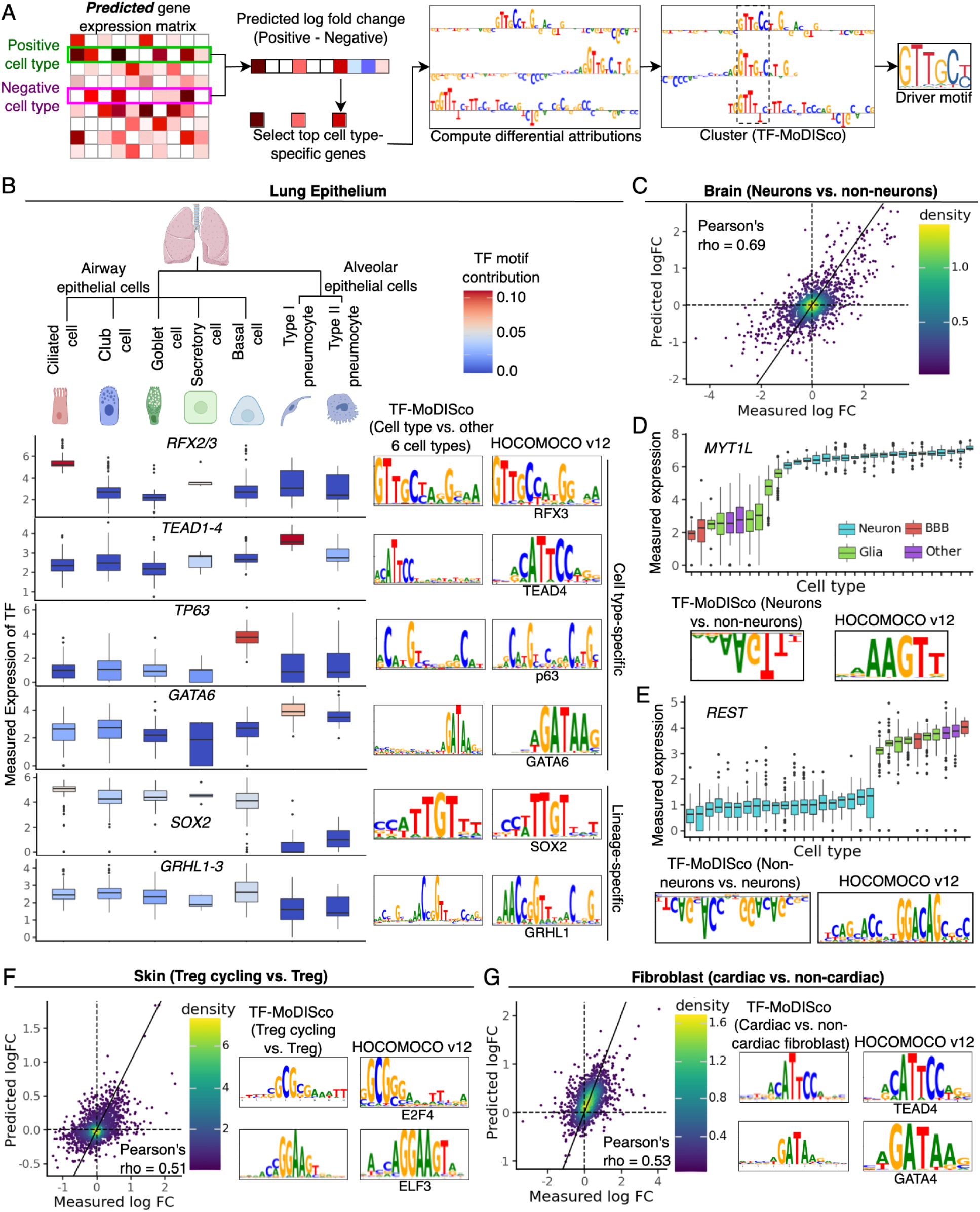
**A)** Schematic showing the method used to identify transcription factor (TF) motifs driving cell type identity based on Decima’s attributions. **B)** Box plots showing the measured expression of the six TFs with the most cell type- or lineage- specific contribution weights across seven epithelial cell types in the human lung. Box plots are colored based on the contribution weight for the corresponding motif in each cell type. For each of the six TFs, the corresponding motif identified by TF-MoDISco is shown on the right. **C)** Scatterplot showing the measured and predicted log fold change in expression of 1,811 test set genes in pseudobulks representing neurons vs. pseudobulks representing other cell types in the brain. **D)** Box plots showing the measured expression of the MYT1L gene in different cell types in the brain, along with the MYT1L motif found by TF-MoDISco based on Decima’s differential attributions (attributions of the difference in predicted expression between neurons and non-neurons). **E)** Box plots showing the measured expression of the REST gene in different cell types in the brain, along with the REST motif found by TF-MoDISco based on Decima’s differential attributions (attributions of the difference in predicted expression between non-neurons and neurons). In D) and E), cell types are colored by their functional categories. BBB = Blood-brain barrier. **F)** Scatter plot showing the measured and predicted log fold change in expression of 1,811 test set genes in cycling regulatory T cells (Treg cycling) vs. non-cycling regulatory T cells (Treg) in skin, along with the top two motifs identified by TF-MoDISco based on differential attribution scores for genes upregulated in Treg cycling vs. Treg. **G)** Scatterplot showing the measured and predicted log fold change in expression of 1,811 test set genes in cardiac fibroblasts vs. fibroblasts in other tissues, along with the top two motifs identified by TF-MoDISco based on differential attribution scores for genes upregulated in cardiac fibroblasts. All TF-MoDISco motifs are accompanied by the most similar motif in the HOCOMOCO v12 database, identified using Tomtom (Gupta et al. 2007).

We performed this analysis on seven epithelial cell types in the human lung, calculating differential attributions for each cell type using the remaining six as ‘negative’ cell types. After clustering with TF-MoDISco and assigning specificity contribution scores to motifs, we identified the motifs that had high contribution scores in only one of the seven cell types, or in a single lineage. The results revealed strong and specific contribution scores for motifs of known lineage-determining TFs (**Supplementary Fig. 6**). Specifically, the top four cell type-specific motifs were: RFX motifs in ciliated cells (Piasecki, Burghoorn, and Swoboda 2010), TEAD motifs in Type I pneumocytes (Little et al. 2021), p63 motifs in basal cells (Daniely et al. 2004), and GATA6 in type I pneumocytes (Yang et al. 2002). We also identified two motifs which contribute specifically to the airway epithelial lineage of cells: SOX2 (Shiraishi et al. 2024) and a motif corresponding to the GRHL family (Kersbergen et al. 2018). In all cases, the role of the TF was further supported by the elevated expression of the TF itself in the corresponding cell type **(Fig. 4B)**.

Since Decima predicted the difference in gene expression between neurons and non-neuronal cells in the brain with high accuracy **(Fig. 4C)**, we tested whether it could identify motifs driving a distinct neuronal identity. TF-MoDISco on differential attribution scores between neuronal and non-neuronal cell types in the brain revealed as the top result a motif with strong negative contributions - i.e. the motif decreases the expression of a gene in neurons relative to non-neuronal cells. This corresponded to the MYT1L transcription factor binding motif **(Fig. 4D)**. Indeed, MYT1L is a pan-neuronal TF **(Fig. 4D)** that establishes neuronal identity by repressing gene programs for other cell types (Mall et al. 2017). In contrast, TF-MoDISco on differential attribution scores between non-neurons and neurons identified a negative contribution for the REST motif - i.e. the REST motif decreases predicted gene expression in non-neurons relative to neurons; **Fig. 4E**). REST is known to repress neuronal genes in non-neuronal cell types (Qureshi, Gokhan, and Mehler 2010). These examples show that Decima’s attributions can identify lineage-specific repressors in addition to activators.

Having found that Decima successfully predicts differences between cell types, we next examined specific cases to test whether it can predict and interpret the finer differences between the same cell type in different tissues or states. For instance, Decima predicted the differential expression of test set genes in cycling vs. resting regulatory T cells, with a Pearson correlation of 0.51 between measured and predicted log fold-changes in expression (cycling vs. non-cycling; **Fig. 4F**). For each gene upregulated in the cycling state, we computed the differential attribution (Treg cycling vs. Treg) for each input nucleotide. We ran TF-MoDISco on these differential attributions, and the top two results matched motifs for the cell cycle related E2F family of TFs (**Fig. 4F**). Similarly, Decima predicted differences in the expression of test set genes between cardiac fibroblasts and fibroblasts in other tissues, with a Pearson correlation of 0.53 (**Fig. 4G**), and differential attribution scores for genes upregulated in cardiac fibroblasts consistently highlighted motifs for GATA4 and GATA6, which are known signature TFs for cardiac fibroblasts (Eraslan et al. 2022; Masuda, Matsuura, and Shimizu 2023), as well as TEAD1, an essential regulator of cardiac fibroblast activation (Song et al. 2024; Burgos Villar, Liu, and Small 2022).

### 2.4 Decima predicts the impact of non-coding variants at cell type resolution

One of the major use cases for sequence-to-expression models is predicting the impact of non-coding genetic variation. In particular, due to linkage disequilibrium, it is often difficult to identify the causal variant for a particular phenotype among several variants in a locus identified by genome-wide association studies (GWAS), and to link these variants to the causal genes that they influence. In addition, it is often difficult to identify the causal cell type or condition in which a variant acts to influence the phenotype. This makes Decima particularly promising for variant effect prediction, owing to its success at predicting cell type- specific expression.

We first asked whether Decima can prioritize variants that significantly impact gene expression in specific cell types. We tested this using a dataset of fine-mapped single-cell expression quantitative trait loci (sc-eQTLs) in PBMCs based on data from the OneK1K consortium (Yazar et al. 2022) and reprocessed by the eQTL catalog (Kerimov et al. 2021). We extracted all 984 high-confidence (PIP >= 0.9) fine-mapped sc-eQTLs for which the variant fell within Decima’s pre-defined input window for the target gene, and matched each one to 20 negative control variants. We used Decima to predict variant effect scores (log fold change in gene expression caused by the variant) for all sc-eQTLs and controls in the matched cell type **(Fig. 5A; Supplementary File 2)**. In each cell type, we tested whether Decima’s scores distinguished the fine-mapped sc-eQTLs in that cell type from matched controls **(Fig. 5B)**. Since the training set of the original Borzoi model also included RNA-seq data from FACS-sorted PBMCs, we repeated this analysis using Borzoi and found that Decima’s performance exceeded that of Borzoi in 19 of 21 cell types as well as in the whole dataset. It also exceeded the performance of a baseline metric (distance of the variant from the TSS; **Supplementary Figure 7-8**).

**Fig. 5.**
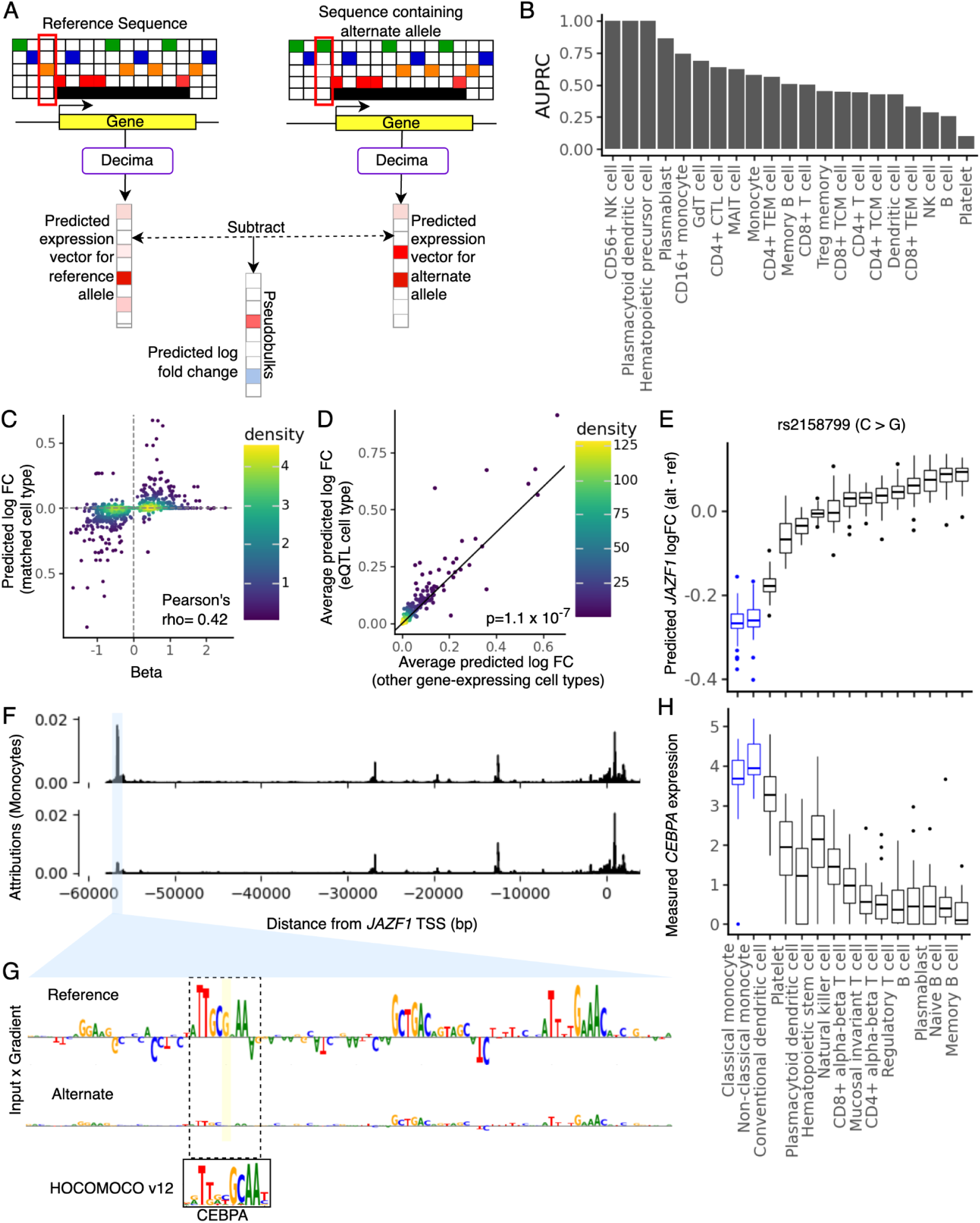
**A)** A schematic showing the procedure for variant effect prediction with Decima **B)** A bar plot showing Decima’s performance in sc-eQTL classification in each of 21 blood cell types in the OneK1K dataset. Performance is measured by the Area Under the Precision Recall Curve (AUPRC) for classification of sc-eQTLs vs. matched control variants in each cell type. **C)** Scatter plot showing Decima’s predicted effect size for high-confidence sc-eQTLs compared to their measured effect size (beta value) in the same cell type. **D)** Decima’s predicted absolute effect size for high-confidence sc-eQTL variants in cell types where the variant is identified as an sc-eQTL with PIP > 0.9, compared to other cell types in which the variant had no significant effect (PIP < 0.1) but its corresponding eGene is still expressed. **E)** Decima’s predicted effect sizes for a monocyte-specific sc-eQTL in various blood cell types. **F)** Decima’s attributions for JAZF1 expression in the sequences containing the reference and alternate allele, showing that the variant (highlighted in blue) reduces the attribution over a distal enhancer. Values plotted represent the absolute input x gradient scores smoothed with a gaussian filter of width 11 bp. **G)** Nucleotide-resolution input x gradient attribution scores computed with respect to predicted expression of JAZF1 in monocytes, for the 160 bp sequence surrounding the reference and alternate alleles of the variant in E-F), highlighting a C/EBP motif disrupted by the variant. **H)** Measured expression of the CEBPA gene over the same cell types shown in E), highlighting its elevated expression in monocytes.

We next asked whether Decima could correctly identify the direction of a variant’s effect on gene expression. For high-confidence (PIP > 0.9) sc-eQTLs, we found a significant positive correlation (Pearson’s rho = 0.42; p-value = 3 x 10^-43^) between the eQTL beta value and Decima’s predicted variant effect in the corresponding cell type **(Fig. 5C)**. Limiting this analysis to the sc-eQTL variants for which Decima predicts any effect (predicted log fold change in expression > 0.01), the correlation coefficient increased to 0.58 (p-value = 2 x 10^-41^), with the direction of effect being predicted correctly in 87% of cases. Thus, for sc-eQTLs that were correctly identified by Decima, Decima also correctly predicts the direction of effect in most cases.

In contrast to previous benchmarks of deep learning-based variant effect prediction that have focused on bulk eQTLs, our sc-eQTL benchmark offers the opportunity to evaluate whether Decima can not only distinguish functional variants but also identify the causal cell type for a non-coding variant. Focusing on 513 variants that were found to be high-confidence sc-eQTLs in only a subset of the 21 cell types in the OneK1K study, we found that Decima’s predicted variant effect scores were higher in the sc-eQTL matched cell type compared to other cell types, even compared to the subset of other cell types in which the target gene is expressed (Wilcoxon test p-value = 1.1 x 10^-7^; **Fig. 5D**).

For instance, in the rs2158799 variant, which is a distal (∼57 kb upstream of the TSS) sc-eQTL for *JAZF1* in monocytes, Decima correctly predicts that the strongest effect of the variant on this gene occurs in monocytes (**Fig. 5E)**. This effect is far stronger than that of the 20 matched negative control variants (**Supplementary Fig. 9**) Decima’s attributions for the surrounding sequence suggest that the variant overlaps with a distal regulatory element that contributes strongly to *JAZF1* expression in monocytes **(Fig. 5F)** but not in other cell types (**Supplementary Fig. 10**). The attributions specifically highlight a C/EBP motif which is disrupted by the alternate allele **(Fig. 5G)**. C/EBP factors are known master regulators of the myeloid lineage including monocytes (Rosenbauer and Tenen 2007), suggesting that this variant acts by disrupting the accessibility of a cell type-specific enhancer.

Given Decima’s success at identifying eQTLs and predicting their impact at cell type-resolution, we asked whether the model could also be useful in interpreting GWAS (Genome-Wide Association Study) variants. This is a significantly harder task for several reasons: (1) Only a subset of GWAS variants alter gene expression; (2) GWAS variants may have weaker effects on expression than high-confidence eQTLs; (3) For GWAS variants, we do not know the causal gene or cell type(s); (4) GWAS variants may impact expression in cell states that are not represented in the dataset.

We selected 837 high-confidence (PIP > 0.9, p-value < 10^-6^) fine-mapped GWAS SNPs for 39 phenotypic traits (**Supplementary Table 2**). Using Decima, we predicted the absolute log fold change in gene expression caused by each variant, on all genes for which the variant fell within Decima’s pre-defined input window. Since we do not know the causal gene, we matched each variant to the gene where it has the strongest predicted impact on expression according to Decima. We further matched each GWAS variant to 10 negative control variants, and predicted their impact on expression of the matched gene using the same procedure (**Supplementary File 3**).

Initially, we compared GWAS variants to controls by assigning each variant a score corresponding to the average predicted absolute log fold change in gene expression caused by the variant, across all cell types. Overall, Decima predicted that GWAS variants caused significantly larger changes in expression than their matched controls **(Fig. 6A)**. Based on these scores, we could classify GWAS variants with an AUPRC of 0.23, outperforming the baseline metric of distance from the variant to the gene TSS **(Fig. 6B**, AUPRC 0.14). 40% of GWAS variants had significantly higher scores (z-test p-value < 0.05) than their matched controls, and 35% had higher scores than all 10 of their matched control variants.

**Fig. 6.**
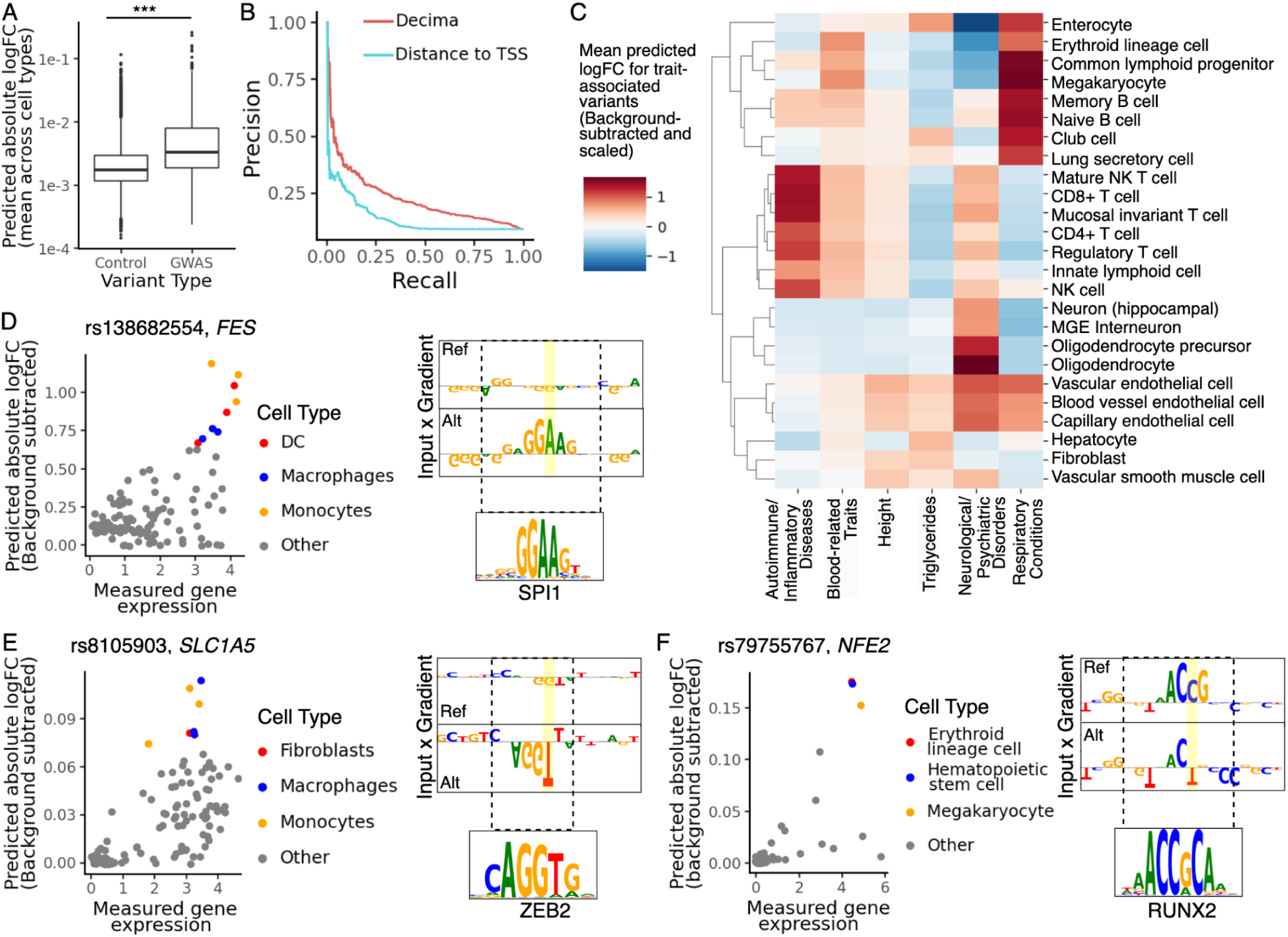
**A)** Box plots showing the distribution of the absolute log fold changes predicted by Decima for 837 high-confidence GWAS variants and 8370 matched negative control variants. **B)** Precision-Recall curves for classification of GWAS variants from matched negative control variants, based on either the predictions shown in A) or on the distance of the variant from the gene transcription start site (TSS). **C)** Heat map showing Decima’s predicted log fold change caused by GWAS variants for 6 traits or categories (x-axis) in selected cell types (y-axis). For each GWAS variant, the average predicted log fold change for all 10 matched negative variants was subtracted from the predicted log fold change for the GWAS variant. The resulting background-subtracted log fold changes across 201 cell types were converted into z-scores. Values in the heatmap are the average z-score for GWAS variants matched to the trait or category. Only GWAS variants that could be distinguished from their matched controls based on Decima’s predictions were included. **D)** Scatter plot showing the measured gene expression (log(CPM)+1) for the FES gene across cell types (x-axis) and Decima’s predicted absolute log fold change in this gene’s expression due to the rs138682554 variant (y-axis). The values on the y-axis represent background-subtracted log fold changes, i.e. the mean absolute log fold change for 10 matched negative control variants was subtracted. Cell types where the variant is predicted to have the strongest impact are highlighted. Decima’s attributions for the sequence surrounding the variant in the highlighted cell types are shown along with the matched motif from HOCOMOCO v12. **E)** Scatter plot showing the measured gene expression (log(CPM)+1) for the SLC1A5 gene across cell types (x-axis) and Decima’s predicted absolute log fold change (background-subtracted) in this gene’s expression due to the rs8105903 variant (y-axis). Cell types where the variant is predicted to have the strongest impact are highlighted. Decima’s attributions for the sequence surrounding the variant in the highlighted cell types are shown along with the matched motif from HOCOMOCO v12. **F)** Scatter plot showing the measured gene expression (log(CPM)+1) for the NFE2 gene across cell types (x-axis) and and Decima’s predicted absolute log fold change (background-subtracted) in this gene’s expression due to the rs79755767 variant (y-axis). Cell types where the variant is predicted to have the strongest impact are highlighted. Decima’s attributions for the sequence surrounding the variant in the highlighted cell types are shown along with the matched motif from HOCOMOCO v12.

It is expected that only a subset of GWAS variants that alter gene expression would be detectable in eQTL studies (Mostafavi et al. 2023), highlighting the necessity of alternate methods to predict variant impact on gene expression. Indeed, only 95 of our 837 selected GWAS variants are also high-confidence eQTLs (11%). As expected, Decima performed especially well at classifying GWAS variants that were also eQTLs, successfully assigning 62% of them significantly higher scores than their matched negatives. This represents a significant enrichment compared to the overall success rate of 39% (Fisher’s exact test p-value 10^-5^) and increases our confidence in Decima’s ability to identify expression-altering GWAS variants. We also attempted to distinguish GWAS variants and controls using RNA or CAGE predictions from Borzoi, and found that, while both models performed similarly, Decima slightly outperformed Borzoi at this task (**Supplementary Table 3**).

A significant advantage of Decima over bulk-trained models is the possibility of linking variants to phenotypes via cell type-specific effects. Taking 334 variants which were successfully distinguished from their matched negative variants (z-test p-value < 0.01) by Decima, we examined the distribution of their predicted effect on expression across cell types. For each variant, we subtracted the mean effect (absolute log fold change in expression) of its 10 matched negatives, then z-scored the resulting differences across cell types. We examined the distribution of these z-scores across cell types for variants matched to different higher-level phenotypic categories **(Fig. 6C)**. Consistent with previous studies, Decima predicted that, on average, variants associated with autoimmune and inflammatory diseases (asthma, eczema, autoimmune disease, Crohn’s disease, inflammatory bowel disease, multiple sclerosis, or lupus) had the strongest effect in T and NK cells (M. J. Zhang et al. 2022). In contrast, variants associated with blood cell related traits (mean corpuscular hemoglobin, RBC count, RBC width, and platelet count) had the strongest predicted effect in erythroid progenitors and megakaryocytes, the precursors of RBCs and platelets respectively. Variants associated with height had the strongest predicted effect in fibroblasts and vascular cells, in line with previous analyses (K. Zhang et al. 2021). Variants associated with blood triglyceride levels were predicted to most strongly alter gene expression in enterocytes and hepatocytes, which are responsible for the absorption and synthesis of triglycerides respectively (Alves-Bezerra and Cohen 2017). Variants associated with neurological and psychiatric disorders (Alzheimer’s disease, Schizophrenia, Bipolar disorder, or neuroticism) were predicted to have their strongest impact in oligodendrocytes, oligodendrocyte precursors, and neurons. Finally, variants associated with respiratory conditions had the strongest predicted effect in lymphoid cells as well as lung epithelial cells such as secretory cells and club cells.

We examined specific variants to ask whether Decima could additionally suggest mechanistic hypotheses underlying functional variants. For example, the rs138682554 variant is associated with hypertension and is also an eQTL in macrophages and monocytes, which are critical for the progression of hypertension (Wenzel 2018). Consistent with the eQTL data, Decima predicts that this variant increases expression of the *FES* gene, particularly in macrophages, monocytes and dendritic cells, and the attributions in these cell types reveal that the variant creates a potential binding site for the myeloid-specific TF SPI1 (**Fig. 6D**). Another example is the rs8105903 variant which is associated with height and is also an eQTL in fibroblasts and monocytes. Consistent with the eQTL data, Decima predicts that this variant decreases expression of *SLC1A5*, particularly in fibroblasts, monocytes, and macrophages, and the attributions in these cell types reveal that the variant creates a potential binding site for the repressive TF ZEB2 (**Fig. 6E**).

Given Decima’s success at predicting cell types consistent with disease biology and eQTL studies, we hypothesize that the model can be a valuable resource to propose cell type-specific mechanisms for the large fraction of GWAS variants that have no associated cell type-specific eQTL. For example, Decima predicted that the rs79755767 variant, associated with RBC width and platelet count, has its strongest effect on the *NFE2* gene in megakaryocytes, erythroid progenitors, and hematopoietic stem cells (**Fig. 6F**). This gene encodes a major regulator of erythroid and megakaryocytic gene expression (“Stimulation of NF-E2 DNA Binding by CREB-Binding Protein (CBP)-Mediated Acetylation” 2001), and Decima’s attributions suggest that the variant disrupts a potential RUNX2 motif (**Fig. 6F**).

These results demonstrate that Decima can identify a subset of non-coding GWAS variants that act by mediating significant expression changes, including in contexts that may not be captured by eQTL studies. Further, in the cases where it is successful, it can be a valuable tool for revealing the underlying cell type-specific mechanisms of action. It is worth noting that Decima provides a prediction of cell type-specific variant impact that is distinct from the expression level of the target gene. For example, the *FES*, *SLC1A5* and *NFE2* genes are all highly expressed in several unrelated cell types, in which Decima predicts only weak effects on gene expression (**Fig. 6 D-F**).

### 2.5 Decima reveals sequence determinants of disease-specific cell states

Decima’s training dataset contains pseudobulks representing 82 disease conditions. Overall, the variation in gene expression between diseased and healthy states of the same cell type is much smaller than the variation between healthy cell types. Nevertheless, we tested whether Decima can predict differences in gene expression between matched healthy and disease samples for the same cell type. We selected pairs of matched healthy and disease pseudobulks for the same cell type in the same tissue and study **(Methods)**, and computed the measured and predicted log fold changes in gene expression (disease vs. healthy) for each pair **(Fig. 7A)**. Across all 565 selected disease/healthy pairs, we found an average Pearson correlation of 0.24 between the measured and predicted log fold changes for test set genes **(Fig. 7B)**.

**Fig. 7.**
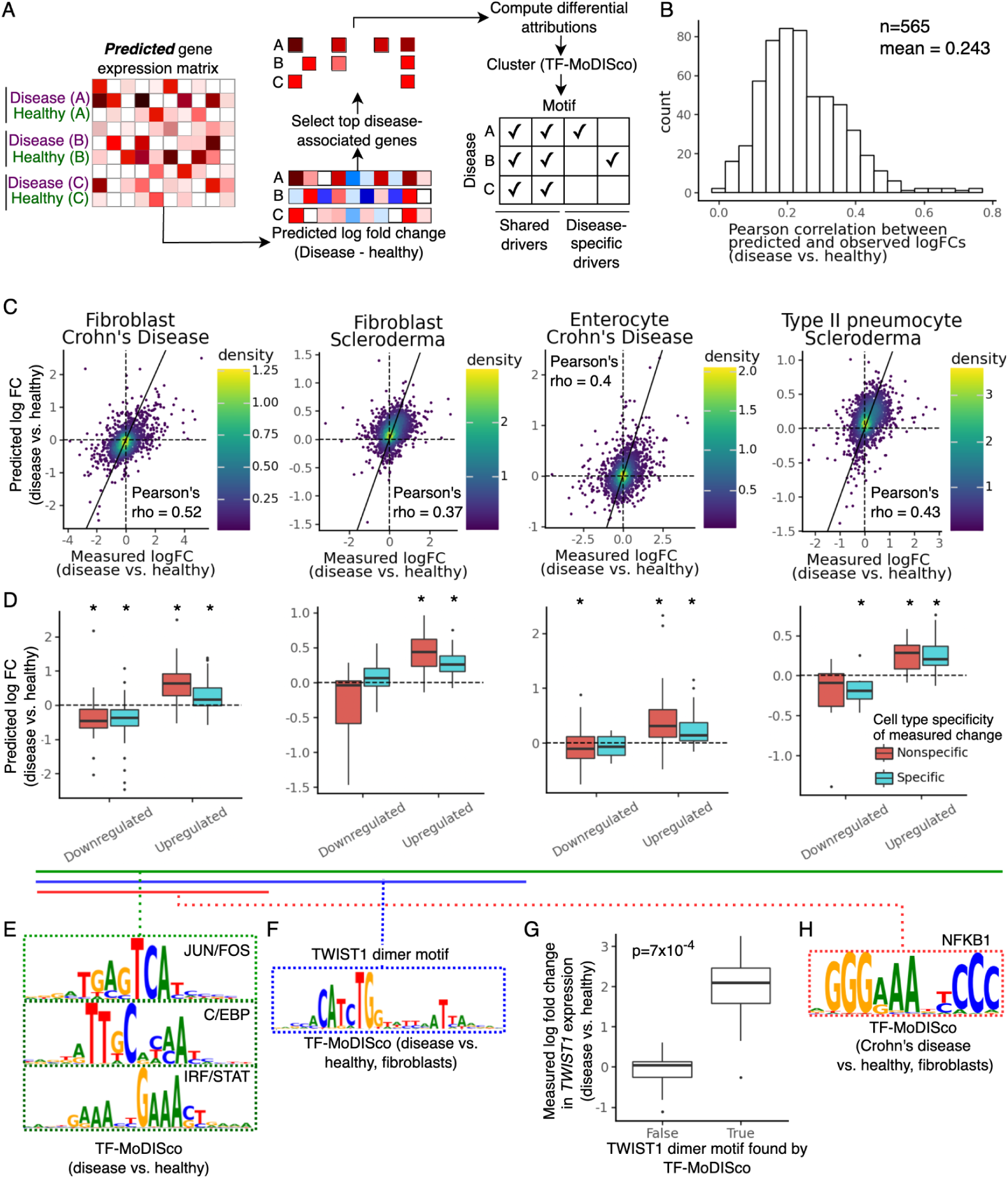
**A)** Schematic showing the method used to identify both shared and disease-specific driver TFs for various cell types and diseases based on Decima’s attributions. **B)** Histogram of Pearson correlations between measured and predicted log fold changes between matched healthy and disease pseudobulks for the same cell type, calculated over 1,811 test set genes **C)** Scatter plot showing the measured and predicted log fold changes for 1,811 test genes between matched healthy and disease pseudobulks, for four cell type / disease combinations **D)** Boxplots showing Decima’s predicted log fold changes for genes that are upregulated or downregulated (measured absolute log fold change >= 1) in disease. Genes are split by whether they are affected by the disease (measured absolute log fold change >= 1) specifically in the cell type of interest, or in multiple cell types. Asterisks indicate groups in which the mean predicted log fold change is significantly higher or lower than 0 (Wilcoxon test p-value < 0.05). **E)** Motifs found by TF-MoDISco based on Decima’s differential attribution scores (disease vs. matched healthy pseudobulks) across multiple cell types and diseases, including all four cases shown in C) and D), representing a shared inflammatory signature **F)** Dimeric motif identified by TF-MoDISco in attributions for fibroblasts in multiple fibrotic diseases, including both fibroblast examples shown in C). **G)** Measured log fold change (disease vs. healthy) in TWIST1 expression in various diseases and cell types, separated by whether the TWIST1 dimeric motif is found by TF-MoDISco based on the disease vs. healthy differential attribution scores in that disease and cell type. **H)** NF-kB motif identified by TF-MoDISco in attributions for fibroblasts in Crohn’s Disease.

For example, Decima predicts the change in expression of test set genes between fibroblasts in the ileum of Crohn’s disease patients compared to fibroblasts in the healthy ileum, with a Pearson correlation of 0.52 **(Fig. 7C)**. Decima correctly predicts the direction of effect for genes that are upregulated (log fold change > 1) or downregulated (log fold change < -1) by Crohn’s disease in fibroblasts, both in cases where the gene is affected by disease specifically in fibroblasts, and in cases where the expression of the gene is altered by disease across multiple cell types in the ileum **(Fig. 7D)**. Similarly, we show that Decima predicts the change in expression of test set genes between enterocytes in the ileum of Crohn’s disease patients compared to the healthy ileum, fibroblasts in the lung of Scleroderma patients and fibroblasts in healthy lungs, and between type II pneumocytes in the lungs of Scleroderma patients and type II pneumocytes in healthy lungs **(Fig. 7 C-D)**.

Given these successful predictions, it follows that Decima can potentially be a powerful tool to identify sequence features and motifs that drive cell type-specific disease responses. Relatedly, we also asked whether Decima can identify shared factors contributing to multiple diseases. To identify TFs that potentially drive differences in gene expression between matched disease and healthy cell populations, we performed differential attribution analysis of genes predicted to be upregulated in disease followed by TF-MoDISco clustering to identify motifs with high contributions **(Fig. 7A)**. In many of the diseases analyzed (specifically scleroderma, atopic dermatitis, Crohn’s disease, idiopathic pulmonary fibrosis, ulcerative colitis, and myocardial infarction), this procedure highlighted motifs corresponding to TF families known to regulate inflammatory pathways, such as JUN/FOS, STAT/IRF and C/EBP **(Fig. 7E; Supplementary Fig. 11)**. In contrast, the contribution of these motifs was lower in chronic kidney disease, which is expected to have a weaker inflammatory contribution.

Beyond this general inflammatory signature, we observed a dimeric motif **(Fig. 7F)** in fibroblasts - but not other cell types - particularly in fibrotic diseases, including Crohn’s disease, idiopathic pulmonary fibrosis, lung scleroderma, as well as COVID-19 in the lung parenchyma. This dimeric motif exhibits a very high-fidelity match to a ChIP-seq motif previously reported for the TF TWIST1 (TWST1.H12CORE.0.P.B) and represents the cooperative binding of TWIST1, together with a homeodomain factor (Kim et al. 2024). We observed a significant association between the presence of this motif in attributions and the upregulation of *TWIST1* gene expression in fibroblasts in specific diseases (Mann-Whitney U-test p-value=7 x 10^-4^ ; **Fig. 7G; Supplementary Fig. 12**). It was previously reported that TWIST1, together with the homeodomain factor PRRX1 which may form the partner in the dimeric motif, controls fibroblast activation in wound healing and in pathological fibrotic conditions (Yeo et al. 2018). Finally, we found an NFκB motif in fibroblasts specifically in Crohn’s disease **(Fig. 7H)**, consistent with reports of NFκB activation in the gut of patients with inflammatory bowel diseases, particularly Crohn’s Disease (Schreiber, Nikolaus, and Hampe 1998; Han et al. 2017).

Our analyses demonstrate that Decima has learned known features of disease biology and can potentially be applied to interpret scRNA-seq data from disease conditions.

### 2.6 Decima can design cell type- and disease-biased regulatory elements

Recent advances in genomic sequence modeling have demonstrated the potential of trained oracle models to generate sequences that drive tissue-specific expression (Taskiran et al. 2024; de Almeida et al. 2024; Gosai et al. 2023; Lal et al. 2024). These model-designed sequences have been shown to drive specific and experimentally validated expression in human cell lines, as well as across various tissues in *Drosophila*. However, a major gap remains: current approaches lack the resolution to tune gene expression at the cell-type level, which is essential for applications such as Adeno-Associated Virus (AAV)-mediated gene therapy (D. Wang, Tai, and Gao 2019). Delivering vectors that target specific cell types in diseased tissues while avoiding healthy cells could also open new therapeutic avenues.

We explored the application of Decima to evolve regulatory elements that are both cell-type and disease-enriched. As a proof of concept, we targeted fibroblasts, which are implicated in the inflammatory response in Crohn’s disease. Fibroblasts contribute to tissue remodeling, fibrosis, and the inflammatory response in Crohn’s disease, making them a relevant target for treatment. We first generated a synthetic construct composed of a random 200bp starting sequence paired with a cargo gene sequence to score expression (**Fig. 8A, Methods**). In order to predict the potency of the regulatory sequence with Decima, this construct was inserted into a genomic safe harbor locus *in silico*. The starting sequence was predicted to have no expression in any cell type, providing a starting point for the design process (**Fig. 8B**).

**Fig. 8.**
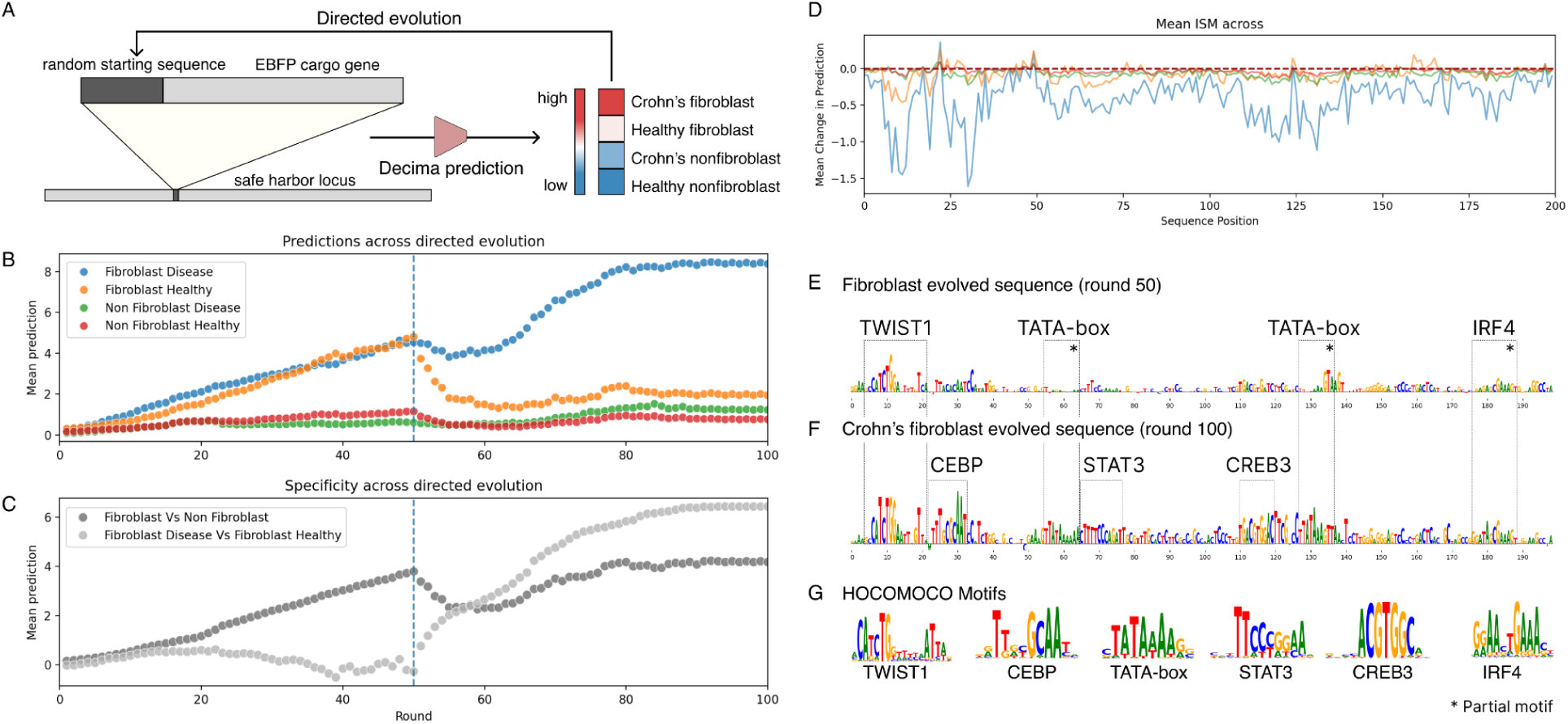
Design of a fibroblast-specific regulatory element in the context of Crohn’s disease using directed evolution. **A**) A schematic showing promoter design through directed evolution with Decima **B)** Predicted expression of the cargo gene across healthy and diseased fibroblast and non-fibroblast cells over 100 rounds of directed evolution. **C)** Predicted specificity of cargo gene expression in fibroblasts and disease fibroblasts, which were optimized in design in rounds 0-50 for cell-type specificity and 50-100 for disease-state specificity. **D)** In silico mutagenesis (ISM) of the synthetic regulatory element reveals key sequence motifs whose perturbation is predicted to uniquely affect expression of the cargo gene in fibroblasts. **E,F)** ISM with respect to disease fibroblast expression identifies key motifs generated in the design process for the fibroblast (top) and disease-fibroblast (bottom) evolved sequences, including a TWIST1, C/EBP, and IRF motif, which are implicated in fibroblast-specific and immune-specific regulation. **G)** These motifs match HOCOMOCO v12 motifs.

To achieve cell type-specific expression, we designed a regulatory element to drive expression specifically in fibroblasts relative to other gut associated cell types such as endothelial cells, macrophages, B cells, and T cells. We performed 50 rounds of directed evolution, maximizing expression in fibroblasts while minimizing expression in the outgroup cell types. After directed evolution, the resulting sequence was predicted to drive robust expression specific to fibroblasts and contained a fibroblast-implicated TWIST1 motif, demonstrating the ability of our approach to generate regulatory elements with cell type-specific expression within a complex tissue environment. We then further evolved this regulatory element to differentiate between fibroblasts from Crohn’s disease patients and those from healthy individuals. We performed an additional 50 rounds of directed evolution, optimizing expression in Crohn’s-derived fibroblasts relative to healthy fibroblasts. The final evolved sequence demonstrated a 6.84-fold higher predicted expression in disease-associated fibroblasts compared to other gut cell types and a 4.31-fold higher predicted expression compared to healthy fibroblasts (**Fig. 8C)**.

To understand the mechanism underlying disease specificity, we applied *in silico* mutagenesis (ISM) to the final evolved sequence, measuring the sensitivity of the predicted cell type- and disease-enriched expression to sequence perturbations. Several motifs are predicted to reduce expression of the cargo gene when perturbed, specifically in Crohn’s-derived fibroblasts (**Fig. 8D**). Among these motifs, we identified two TBP motifs (TATA-binding protein), essential in transcription initiation (**Fig. 8E-G**). This suggests that the model designed an effective and standalone promoter (Ponjavic et al. 2006). The TWIST1 motif was consistent in both fibroblast-specific and disease-fibroblast-specific evolved sequences, aligning with previous findings implicating this motif in fibroblast-specific expression. As we evolve the sequence toward disease-fibroblasts, we also observe the completion or installation of C/EBP, STAT3, CREB3, and IRF motifs. Several of these motifs were previously identified as associated with inflammatory response, indicating that the model is implanting motifs it has observed to drive inflammatory and cell-type specific expression. This proof-of-concept study demonstrates the potential of Decima to design highly specific regulatory elements that can distinguish not only between cell-types, but also between disease states.

## 3. Discussion

Sequence-to-function models have become pivotal in understanding how the genome sequence is interpreted and executed in various cellular contexts. These models are essential for generating mechanistic hypotheses for non-coding variants and designing synthetic regulatory sequences. Although recent models have expanded their biological coverage to include various experiments and tissues, their reliance on bulk samples from healthy individuals and cell lines limits the investigation of regulatory circuits specific to cell types or diseases. Single-cell RNA sequencing (scRNA-seq), for which substantial datasets are already publicly available, can overcome these limitations by providing extensive coverage of diverse cell states and diseases, thereby enriching our understanding of complex regulatory rules.

However, the field of single-cell genomics faces challenges in inferring regulatory activity in individual cell types and states when chromatin accessibility data is unavailable. This limitation also significantly restricts the generation of hypotheses regarding the regulatory elements driving differential expression between cell types and cellular states. Decima bridges this gap by applying the utility of sequence-to-function models to single-cell datasets, enabling regulatory interpretation in various cell states and diseases, sc-eQTL effect predictions, and the design of regulatory sequences with cell type- and disease-enriched activity. Our findings show that Decima can learn regulatory rules related to cell type specificity, tissue residency, and disease-dependent cell states by replicating known regulatory syntax from the literature. We believe that the scRNA-seq community will utilize Decima to add an extra layer of regulatory insight to the results generated by current scRNA-seq analyses, thereby enhancing the comprehension of biological phenomena.

One caveat of our approach is the potential for experimental batch effects, which might introduce spurious regulatory sequences in the downstream analyses. Modeling regulatory logic in cell populations that exist in a continuum and are difficult to categorize into distinct cell types also poses a challenge in the current formulation, where we collapse cells belonging to the same discrete group into a single pseudobulk. Further, as we are primarily focused on predicting expression differences between cell types rather than between individuals, we also collapsed cells from different individuals in the same study into the same pseudobulk.

Caveats of most existing sequence-to-expression models, including those using bulk expression data, also hold here: by defining instances as the sequence surrounding a gene and its expression across pseudobulks, Decima captures *cis*-regulatory mechanisms. Models that incorporate broader cellular state information, for example defined by some appropriate representation of the full transcriptome, or by subset of appropriately selected genes, would be useful extensions that could capture some *trans*-regulation effect. Moreover, the output space of this model is strictly defined by the cell-type, tissue, disease combinations in our training set and thus the model is not designed to transfer to unseen cellular contexts.

Additionally, there is relatively low concordance between the disease versus healthy changes in expression measured in multiple studies of the same disease, presenting further complications in modeling the regulatory biology of diseases. The predictability of disease differential expression, arguably the most difficult task in our study, depends heavily on the model’s performance for the specific genes and cell types involved in the disease, as well as the regulatory nature of the disease itself. Nonetheless, for diseases with clear disease-specific or enriched cell states, identifying regulatory drivers is not markedly different from identifying cell type-specific regulatory elements, highlighting the potential of Decima to tackle regulatory challenges in disease biology.

As future work, we plan to build large atlases of predicted functional variants, including rare variants not captured in eQTL studies, in the context of hundreds of cell types. This will allow us to explore the colocalization of these variants with GWAS variants to link disease-associated variants to specific cell types and to identify transcription factors and other regulatory mechanisms and pathways that drive cell type-specific changes in activity in the context of disease. Further, we aim to extend our approach to design more complex regulatory elements that activate therapeutic expression in diseased cells, but turn off when the cells return to a healthy state, opening up a dynamically disease-conditioned approach to gene therapy.

## 4. Methods

### 4.1 Processing sc/snRNA-seq datasets

#### 4.1.1 Data sources

Publicly available sc/snRNA-seq count matrices were downloaded from the following sources:

- SCimilarity (sc/snRNA-seq): Individual datasets were downloaded and prepared as described in (Heimberg et al. 2023).
- Brain cell atlas (snRNA-seq): https://www.braincellatlas.org/dataSet
- Human skin atlas (scRNA-seq): https://singlecell.broadinstitute.org/single_cell/study/SCP2738
- Human retina atlas (snRNA-seq): https://cellxgene.cziscience.com/collections/4c6eaf5c-6d57-4c76-b1e9-60df8c655f1e

#### 4.1.2 Aggregation and filtering of pseudobulks

We summed the count vectors for all cells corresponding to the same cell type, tissue, disease and study into a single pseudobulk, producing a pseudobulk x gene count matrix using the scanpy.get.aggregate(adata, metadata_columns, func=’sum’) function in scanpy version 1.10.2 where ‘adata’ represents the cell-level count matrix and ‘metadata_columns’ denotes the metadata columns identifying each pseudobulk. Note that in this process, cells from different individuals in the same study are summed into a single pseudobulk.

We dropped pseudobulks corresponding to cell lines, cancers, organoids, or unannotated cell types. From the SCimilarity dataset, we also dropped pseudobulks corresponding to samples from the brain, skin, and retina, as these tissues were better represented and annotated in the other atlases. Since cell types in the SCimilarity dataset were automatically annotated using an embedding-based nearest-neighbors approach, we also manually examined the cell type annotations and removed cell types that were likely misannotated (e.g. cell type annotations inconsistent with the tissue). Finally, we removed extremely low quality / sparse pseudobulks that were composed of fewer than 50 cells and which were also in the lowest 10% of pseudobulks in terms of number of genes measured and total counts.

#### 4.1.3 Gene filtering and annotation

We dropped genes and pseudobulks for which > 33% of expression values were missing. We then annotated genes with their start and end coordinates in the hg38 genome using genome annotation files obtained from CellRanger (https://www.10xgenomics.com/support/software/cell-ranger/latest) and the NCBI gene database (https://www.ncbi.nlm.nih.gov/gene). We removed all genes except those in autosomes or Chromosome X.

#### 4.1.4 Normalization

We normalized each pseudobulk such that the total counts across all genes equaled 1 million, followed by applying a log(1+x) transform to all counts. Thus, the final values in the matrix represent log(1+CPM) values.

#### 4.1.5 Creating inputs for Decima

For each gene, we created a 524,288 bp genomic interval starting at 163,840 upstream of the TSS and extending through the gene body. The gene start was placed at this position because the Borzoi architecture lacks information to make fully accurate predictions for bases closer to the input boundary than this (Linder et al. 2023)In cases where the interval extended near or beyond the end of the chromosome, the interval was shifted to be at least 10 kb away from the chromosome end. For each interval, we extracted the corresponding sequence from the hg38 genome assembly and dropped intervals where > 40% of the bases were Ns. In all but 643 of the remaining 18,457 genes, the interval extended beyond the gene end.

To create inputs for Decima, we took the genomic sequence for the interval corresponding to each gene, and reverse complemented the sequence for genes on the negative strand. We one-hot encoded each sequence to obtain a matrix of dimensions (4 x 524,288). To this, we added a fifth row containing a binary mask that highlighted the location of the gene of interest, i.e. the value of the mask was 1 for all positions between the gene start and end, and 0 for all other positions. Combining the one-hot encoded sequence and the gene mask resulted in a matrix of shape (5 x 524,288) representing each gene.

Since we initialized Decima using the weights of the trained Borzoi model, we split genes into training, validation and test sets based on their overlap with the genomic regions used for these sets in Borzoi. As a result, Decima’s test set of 1,811 genes do not overlap with any genomic region used for training or validation of Borzoi. For genes in the training set, we also extracted and added the one-hot encoded sequence for 5000 additional bases at the start and end of the interval. This was done to allow data augmentation by shifting the input window during training, as described below.

### 4.2 Decima architecture

We downloaded the PyTorch version of the Borzoi model (Linder et al. 2023) from gReLU (Lal et al. 2024). We removed the terminal linear layer of this model and added an average pooling layer along the length axis, followed by a linear layer that takes in a vector of size 1920 for each input and returns a vector of size 8,848.

Since Decima takes inputs of size (5 x 524,288) whereas Borzoi takes inputs of size (4, 524,288), we also added an additional input channel to the first convolutional layer of the model.

### 4.3 Training Decima

Decima’s loss function is based on the two-component loss function used in Borzoi (Linder et al. 2023). However, it is applied across the pseudobulk axis for a single gene rather than within individual pseudobulks. Specifically, the loss function consists of two terms: a Poisson loss applied to the total value of the gene across all pseudobulks, and a multinomial loss function that compares the true and predicted distribution of expression values across pseudobulks for the same gene. During training, we applied a weight of 10^-4^ to the poisson component. This downweighting encourages the model to learn to predict differences in expression of the same gene between pseudobulks.

The entire model was trained for 15 epochs with the Adam optimizer, a learning rate of 3x10^-5^, and a batch size of 4 with gradient accumulation over every 5 batches. We performed data augmentation during training by randomly shifting the input sequence by up to 5,000 bases in either direction along the genome. The validation loss was measured after each epoch and the model with the lowest validation loss was saved. The training procedure was performed on a single NVIDIA A100 GPU and took approximately one day. We repeated this 4 times, each time using a different replicate of the Borzoi model (Linder et al. 2023).

### 4.4 Classification of cell type-specific genes

To determine cell-type specific genes, we followed a similar procedure as (Eraslan et al. 2022) at the pseudobulk level **(Fig. 2A)**. First, we z-score normalized the expression of each gene across pseudobulks. We then computed, for each gene, its mean z-score per cell type by averaging across all pseudobulks annotated as corresponding to this cell type. For each cell type, we then labeled the genes which had mean z-score greater than 1 in that cell type, as being specific to that cell-type.

We repeated the z-score normalization and averaging for the predicted expression values and used the ranking implied by the predicted mean z-scores to classify whether a gene was specific for a particular cell-type (i.e. whether it had an observed mean z-score > 1). We note that the number of specific genes (and thus the implied class balance) will vary by cell type. For this reason, we report AUROC as the main metric, as the AUROC is insensitive to differences in class-balance and the performance of a random classifier will be 0.5 in every cell-type. For the purpose of computing the AUROC metric, only test set genes were considered.

### 4.5 Comparing attributions between sequence regions

Coordinates of all annotated cis-regulatory elements (CREs) in the hg38 genome were downloaded from the ENCODE database (ENCODE Project Consortium et al. 2020). Coordinates of exons for all genes were taken from the genome annotation file downloaded from CellRanger (https://www.10xgenomics.com/support/software/cell-ranger/latest).

For each gene in the test set, we averaged the predicted expression over all pseudobulks where the gene was expressed above a threshold (log(CPM+1) > 1.5). We used the Input x gradient method to compute the attribution of this average expression for each nucleotide in the 524,288 bp input sequence. For each gene, we then computed the average of the absolute value of this attribution over all nucleotides overlapping with the annotated exons of the gene, the promoter (+/- 100 bp from the gene TSS), and exon/intron junctions (+/-10bp surrounding each junction). For nucleotides in introns, we computed separate averages for those nucleotides that overlapped with an annotated CRE (‘Intronic CREs) and those that did not. For nucleotides outside the gene body, we divided these into categories based on their distance from the gene body (<1000bp, 1000-10,000bp, 10,000 - 100,000 bp and >=100,000 bp) and further divided each category into nucleotides that overlapped with an annotated CRE and those that did not. We then calculated the average of the absolute value of the Input x gradient attributions over all nucleotides in each category.

### 4.6 Cell type specificity interpretation

To find lineage and cell type-specific motifs in the lung epithelium, we proceeded as follows. First, we computed the cell type specificity of expression for each test set gene using the z-score method as done previously, except that we restricted the underlying gene-expression matrix only to lung / airway-related tissues. For each of the main epithelial cell types (ciliated cell, club cell, secretory cell, goblet cell, respiratory basal cell, type I pneumocyte and type II pneumocyte), we then selected the top 250 most specific genes (i.e. the 250 genes with the highest mean z-score for this cell-type). For each of these genes, we computed the gradient of the difference in gene expression between the cell type for which the gene was specific and the remaining epithelial cell types, with respect to each nucleotide in the input sequence. This gives a ‘differential attribution’ which highlights the parts of the sequence which the model considers relevant for driving cell type-specific expression of genes in each individual epithelial cell type (Lal et al. 2024). We then used TF-MoDISco (Shrikumar et al. 2018) to assemble these attributions into motifs. To limit the considerable runtime of TF-MoDISco, we restricted the clustering to the region +/-10kb around the annotated TSS of the gene, and we restricted the maximum seqlets to 10000. Otherwise, we used default parameters.

After TF-MoDISco clustering, we matched the resulting motifs to HOCOMOCO v12 motifs (Kulakovskiy et al. 2018) using Tomtom (Gupta et al. 2007). Since many TFs share similar motifs, we clustered the HOCOMOCO motifs using the GimmeMotifs software (Bruse and van Heeringen 2018). We then associated each motif hit from TF-MoDISco to the respective motif cluster and aggregated hits for the same cluster. We excluded some motifs, specifically those corresponding to certain Zinc Finger TFs, because their low information content can lead to highly significant but low-quality Tomtom matches.

For each detected motif cluster, we computed an aggregate “specificity contribution score” based on TF-MoDISco as follows: we counted the total number of positive seqlets which could be matched to any non-excluded motif with q-value < 0.05, and then computed the ratio of the number of seqlets found for this motif to the total number of seqlets. If a motif cluster was not detected in a particular cell-type, the corresponding score is thus zero. We excluded the negative seqlets and negative motifs since in such a differential analysis, they may correspond to motifs of other cell-types rather than cell-type-specific repressors. We selected the 4 motifs where this score was most strongly enriched for a specific cell type or lineage to highlight.

To find motifs which differentiate individual cell types, cell states or the same cell type in different tissues, we proceeded as follows. Firstly, we selected the tracks corresponding to the ‘positive’ condition (for the fibroblast example, this would be all fibroblast pseudobulks with organ = ‘heart’) and those corresponding to the ‘negative’ condition (all other fibroblast tracks). On the test set genes, we computed the correlation between predicted and observed fold change between the positive and negative pseudobulks. From all test genes, we then selected the 300 genes which the model predicts to be most upregulated in the ‘positive’ condition and computed the gradient of the difference in predicted expression between the ‘positive’ and ‘negative’ condition. We then performed TF-MoDISco clustering as above. Highlighted motifs were found by manual inspection of the results.

Note that for these analyses, all predictions and attributions were averaged across all four model replicates. Additionally, we subtracted the mean attribution across nucleotides of the gradient, as in (Majdandzic, Rajesh, and Koo 2023).

### 4.7 Processing sc-eQTL data

We downloaded the reprocessed OneK1K (Yazar et al. 2022; Kerimov et al. 2021) sc-eQTL data from the EBI eQTL [catalogue](https://www.ebi.ac.uk/eqtl/) FTP server https://ftp.ebi.ac.uk/pub/databases/spot/eQTL/susie/QTS000038/. We filtered out all variants which cannot be scored by Decima because they are too far from their target gene, as well as variants in non-standard chromosomes and ENCODE blacklist regions. Additionally, because indels can change the length of the underlying sequence making comparisons more difficult, we chose to only analyze single nucleotide variants.

For each cell type in the OneK1K data, we then collected the SuSiE (G. Wang et al. 2020) finemapped eQTL variants which achieved posterior inclusion probability (PIP) greater than 0.9 for some gene. These variants are very likely to be causally impacting the expression of their target gene and thus form the positive set for our eQTL analyses. For each pair of high-PIP eQTL variant and target gene, we then extracted 20 variants which were tested for the same gene but did not achieve statistical significance, did not form part of any SuSiE credible set for any gene in any cell type and have allele frequency > 5%. Where more than 20 such variants were available, we prioritized the ones closest to the TSS of the target gene of the corresponding high-PIP variant (as many causal variants tend to be close to the TSS). These variants formed the negative set, since the data provides no evidence that they have a measurable impact on gene expression. Accordingly, this procedure produced a class-balance of 1:20.

### 4.8 sc-eQTL Variant effect prediction

We matched Decima cell types to OneK1K cell-types as shown in **Supplementary Table 3**. Note that we restrict the matching only to pseudobulks coming from tissue “blood”, i.e. we do not include, for example, liver macrophages. Gamma-delta and double-negative T-cells are not matched, since they were not annotated in our training data. For these cell types, we compute the average predicted variant effect across all blood cell types.

To compute the predicted variant effect for Decima, we proceeded as follows. For each variant-gene pair, we first took as reference sequence the genomic interval around the gene as defined previously (4.1.5). To construct the alternate sequence, we replaced the reference allele with the eQTL alternate allele. We then computed Decima predictions for both the reference and alternate sequence and recorded the difference in prediction as the variant effect **(Fig. 5A)**. This represents a log fold change as Decima’s predicted values are in log1p scale. Note that the model will predict one such variant effect per pseudobulk. To get a consolidated score, we averaged the effect across all pseudobulks which matched the sc-eQTL cell-type of interest, using the matching defined above.

For Borzoi, we computed two different matchings. Firstly, we used the “GTEx blood track”, which measures how well bulk tissue-derived variant effect predictions correspond to deconvolved single-cell eQTL data. Secondly, we manually matched Borzoi tracks to the individual cell types. In this matching we prioritized RNA tracks wherever available, but resorted to CAGE tracks if they more closely matched the cell type of interest. If no matching track was found, we manually selected the most similar cell-type or group of cell-types (e.g. for mucosal invariant T-cells, we matched to the average across all T-cell tracks).

To compute the variant effect for Borzoi, we proceeded as follows. For each gene-variant pair, we provided to Borzoi the same reference/alternate sequences as we provided to Decima. As done in the Borzoi paper (Linder et al. 2023), we removed the cropping layer and recorded the output of all 16,384 bins for all Borzoi expression tracks. For RNA tracks, we undid the Borzoi “squashed-scale” normalization, then summed predictions across all bins which intersected the Decima gene mask, which covers all annotated transcripts of the gene of interest. This collapses the Borzoi prediction to one gene expression score per track, an approximation which is necessary for Borzoi, but not Decima, because Borzoi predicts a gene-agnostic expression profile along the input sequence which needs to be subsequently matched to individual genes. Finally we applied a log transform and computed the log fold change between the predictions for the alternative and reference alleles. For CAGE tracks, we followed a similar procedure except that we summed only +/-10 bins around the beginning of the gene mask, which should cover the TSS and core promoter. As with Decima, when more than one track was matched to the corresponding sc-eQTL cell type, we averaged the predicted variant effect across these tracks. For both Borzoi and Decima, predictions were averaged over all four model replicates.

To test the cell type specificity of Decima eQTL predictions, we first identified cell type specific variants. For this, we selected fine-mapped variants which achieved a PIP > 0.9 for a particular expression gene (eGene) in some cell-type. For each such variant-gene pair, we denoted all the cell types where the variant achieved PIP > 0.9 as “on-target” cell types. All cell types where the variant achieved PIP < 0.1 were considered “off-target” cell types. Cell-types where the variant had an intermediate PIP were excluded. Moreover, we excluded off-target cell-types where the gene was not expressed, i.e. had an average log1p expression of less than 1.5. This threshold was chosen as it separates the two modes of the distribution of average expression (across pseudobulks) of all genes. We then matched the on-target and off-target cell-types to Decima pseudobulks using the same procedure as described above. In some cases the resolution of the OneK1K data exceeds that of Decima, e.g. in the case of subtypes of CD4 T-cells. In this case, if a variant-gene pair is “on-target” for, e.g., one type of CD4 T-cell, then we considered it on-target for CD4 T-cells altogether. After matching, we first averaged Decima variant effect predictions per cell type (to account for the fact that some cell-types are represented by more pseudobulks than others) and then averaged the predictions for on-target and off-target cell-types respectively.

### 4.9 Processing GWAS data

To identify a set of high probability disease causing variants, we conducted fine mapping using summary statistics from 39 traits that had “gold-standard” gene sets defined by Zhang et al. (M. J. Zhang et al. 2022) (**Supplementary Table 2**). Our fine-mapping pipeline consists of three steps under the assumption of a maximum of 5 causal loci in the 1000000 bases surrounding each lead SNP. In the first step, we used DENTIST (W. Chen et al. 2021) to identify and remove SNPs that are discordant with patterns of LD in a defined reference panel, in this case, 50000 individuals from the 380k UKBiobank reference panel (s3://broad-alkesgroup-ukbb-ld/UKBB_LD/). In the second step, we made use of PolyFun (Weissbrod et al. 2020) to compute functional priors for fine mapping using the the baseline annotations (baselineLF2.2.UKB) provided by Gazal et al (Gazal et al. 2018) and L2-regularized S-LDSC (stratified LD score regression). In the third step, the resulting priors were fed to SuSiE (G. Wang et al. 2020) to generate a credible interval around each lead SNP and to assign posterior inclusion probabilities (PIP) to individual SNPs.

From the fine-mapping results, we selected high-confidence (PIP > 0.9 and p-value < 10^-6^) GWAS SNPs within 100 kb of any annotated TSS. Further, we selected the subset of these SNPs that were annotated as regulatory variants in gnomAD (Karczewski et al. 2020), and did not overlap with ENCODE blacklist regions. These 837 variants formed the positive set for our GWAS analyses.

Since we do not know which gene is affected by these variants, we predicted the effect of each positive variant on all genes for which the variant overlapped with Decima’s prediction interval. We matched each gene to the variant with the highest predicted effect (mean across all output cell types/tracks). For Borzoi, we computed variant effect scores as described above and selected the best gene in two ways; based on the average score over RNA tracks or over CAGE tracks. For both Borzoi and Decima, predictions were averaged over all four model replicates.

For each of these ‘positive’ GWAS variants, we then extracted 10 SNPs which were annotated as regulatory and had MAF > 1% according to gnomAD, did not have PIP > 0.01 for any trait in the fine-mapped GWAS datasets, did not overlap with blacklist regions, and were between 10-120,000 bp from the TSS of the matched gene. Where more than 10 such variants were available, we selected the 10 variants whose distance from the TSS was most similar to that of the GWAS variant. These variant-gene pairs formed the negative set. Variant effect predictions for these variant-gene pairs were made using Decima and Borzoi as described above.

To analyze variant effect predictions at the cell type level, we averaged Decima’s predicted variant effect scores for all tracks corresponding to the same cell type. To identify which high-PIP GWAS variants are also known eQTLs, we used fine-mapped eQTL data provided by Open Targets using the September 2022 release (https://ftp.ebi.ac.uk/pub/databases/opentargets/genetics/22.09/). Specifically, we downloaded all data in the “v2d_credset” analysis directory and filtered the results for type “eqtl” and a posterior probability (PIP) of at least 0.9.

### 4.10 Cell type specificity analysis for GWAS variants

We selected GWAS variants which were predicted to have significantly higher impact (z-test p < 0.01) than their 10 matched control variants according to Decima. Due to the limited number of such variants for individual traits, we grouped variants corresponding to similar traits into general categories. Specifically, Inflammatory Bowel Disease, Crohn’s disease, asthma, eczema, autoimmune disease, lupus, and multiple sclerosis were combined into the category ‘Autoimmune and Inflammatory Diseases’. Mean corpuscular hemoglobin, RBC width, RBC count, and platelet count were combined into ‘Blood-related traits’.

For each remaining variant, we computed the difference between Decima’s absolute predicted log-fold change (VEP score) for the variant and its for its 10 matched negative control variants, in each cell type. We then converted these into z-scores and averaged the per-cell type z-scores for all variants corresponding to the same trait (or category).

### 4.11 Disease interpretation

We first selected all pseudobulks from disease samples in the data which could be matched to a healthy control pseudobulk using the following criteria:

- A corresponding healthy pseudobulk from the same tissue, cell type and study was available.
- Both the disease and the corresponding healthy pseudobulk were supported by at least 500 cells each.
- The absolute log-difference in size factor (sum of expression values for all genes) between the disease and the corresponding healthy pseudobulk was at most 0.5.

This resulted in a robust set of pairs of matched disease and healthy pseudobulks, with each pair corresponding to a unique “quadruplet” of cell type, tissue, study and disease. For these pairs, we then computed the observed and predicted log fold change in gene expression between the disease state and the corresponding healthy pseudobulk **(Fig. 6A)**. Using only test set genes, we computed the correlation between the observed and predicted log fold changes.

For a subset of case studies, selected from the diseases where the model performs best on test-genes, we selected the 300 genes where Decima predicted the highest upregulation in disease and computed the gradients for the differential gene expression in disease vs. healthy states. We then used TF-MoDISco clustering to find motifs, as described above.

### 4.12 Regulatory element design

The synthetic element includes a 200 bp regulatory sequence appended to the EBFP cargo gene (blue emission fluorescent protein). The cargo gene sequence was obtained from the GenBank database (GenBank ID: MH450096.1) (Grajevskaja et al. 2018). The synthetic element was inserted in a genomic safe-harbor locus located at chr22:29316378-29840666 *in silico*. This locus was selected due to its lack of endogenous gene expression. All *in silico* testing was evaluated using this placement location to ensure consistency across iterations. This construct was introduced 5120*32 basepairs (163,840bp) upstream of the gene start location within the 524,288 bp input sequence, which matched the gene location observed by the model during training. At each round of directed evolution, we performed *in silico* mutagenesis across the 200bp regulatory element and selected the single nucleotide mutation that maximized the difference in predicted expression between two target groups.

We selected fibroblast and non-fibroblast pseudobulks from a previously published Crohn’s disease study which formed part of Decima’s training set (Collection ID: 17481d16-ee44-49e5-bcf0-28c0780d8c4a) (Elmentaite et al. 2020). Specifically, we selected 1 healthy fibroblast sample, 1 UC fibroblast sample, 22 non-fibroblast healthy samples, and 22 non-fibroblast UC samples. In the first 50 rounds of evolution, the regulatory elements were optimized to drive expression specifically in fibroblasts relative to non-fibroblast cells, i.e. we maximized the difference in average expression between the 2 fibroblast pseudobulks and all 44 non-fibroblast pseudobulks. During the remaining 50 rounds, we refined the design to enhance expression in Crohn’s-derived fibroblasts compared to healthy fibroblasts, optimizing the sequence to maximize differential expression between the Crohn’s-fibroblast pseudobulk and the healthy fibroblast pseudobulk.

Motifs were identified using the HOCOMOCOv12 database (Kulakovskiy et al. 2018) and gReLU (Lal et al. 2024).

## Software availability

Decima is available at https://github.com/Genentech/decima. The code used to process data, train models, and perform all analyses in this paper is available at https://github.com/Genentech/decima-applications. A Snakemake-based pipeline for GWAS fine-mapping is available upon request.

## Data Availability

Sources for all single-cell and variant datasets used in this study, as well as predictions made by Decima for all genes and variants, are given in the supplementary material. Model weights and supplementary material are available at https://doi.org/10.5281/zenodo.13910841.

## Acknowledgments

We would like to thank the following for their helpful comments: Aviv Regev, Jason Rock, Oana Ursu, Heinrich Jasper, Tim Sterne-Weiler, Xiaosai Yao, Sara Mostafavi, Chris Cox, M. Hasan Celik, Kipper Fletez-Brant, Diana Chang, Nikolas Jorstad, Jane Song.

